# Super-exponential growth and stochastic size dynamics in rod-like bacteria

**DOI:** 10.1101/2022.05.21.492931

**Authors:** Callaghan Cylke, Shiladitya Banerjee

**Affiliations:** Department of Physics, Carnegie Mellon University, Pittsburgh, PA 15213, USA

## Abstract

Proliferating bacterial cells exhibit stochastic growth and size dynamics but the regulation of noise in bacterial growth and morphogenesis remains poorly understood. A quantitative understanding of morphogenetic noise control, and how it changes under different growth conditions, would provide better insights into cell-to-cell variability and intergenerational fluctuations in cell physiology. Using multigenerational growth and width data of single *Escherichia coli* and *Caulobacter crescentus* cells, we deduce the equations governing growth and size dynamics of rod-like bacterial cells. Interestingly, we find that both *E. coli* and *C. crescentus* cells deviate from exponential growth within the cell cycle. In particular, the exponential growth rate increases during the cell cycle, irrespective of nutrient or temperature conditions. We propose a mechanistic model that explains the emergence of super-exponential growth from autocatalytic production of ribosomes, coupled to the rate of cell elongation and surface area synthesis. Using this new model and statistical inference on large datasets, we construct the Langevin equations governing cell size and size dynamics of *E. coli* cells in different growth conditions. The single-cell level model predicts how noise in intragenerational and intergenerational processes regulate variability in cell morphology and generation times, revealing quantitative strategies for cellular resource allocation and morphogenetic noise control in different growth conditions.

## I. INTRODUCTION

Uncovering the quantitative principles of single-cell physiology demands high quality experimental data at the single-cell level with extensive statistics [1]. This information needs to be integrated with quantitative theory to interpret single-cell behaviors, inform the underlying mechanistic models, and direct new experimental research. Recent advances in single-cell imaging and microfluidics have resulted in large amounts of high-quality datasets on the size and shapes of single bacterial cells as they grow and divide [2–7]. These data have revealed many fundamental models and principles underlying single-cell physiology, including the mechanisms of cell size homeostasis and division control [4, 8–13], cell size control and growth physiology [12, 14–16], cell shape control [11, 17–19] and adaptation to environmental changes [20– 23]. While extensive work has been done to characterize cell size regulation and division control at the inter-generational level [12], much less is understood about the dynamics of cell growth within an individual cell cycle. The bacterial cell cycle is composed of complex coupled processes, including DNA replication, cell wall synthesis and constriction, that have to be faithfully coordinated for cells to successfully divide. These processes require dynamic remodelling of the cell envelope and shape, raising questions on how cell growth and size changes are dynamically coupled and how noise in these processes are regulated to ensure morphological stability through cycles of growth and division.

In this article, we develop quantitative theory for the stochastic growth and size dynamics of rod-shaped bacterial cells using multigenerational growth and morphology data of *Escherichia coli* and *Caulobacter crescentus* cells [4, 11]. While there are existing studies using stochastic and deterministic models to describe bacterial growth and division processes [24–29], a common assumption in these models is that bacteria grow exponentially in cell size. Few current models deviate from exponential growth, among both bacterial and eukaryotic cells [30]. Our analysis reveals that single *E. coli* and *C. crescentus* cells elongate faster than an exponential during the cell cycle, challenging existing models of exponential growth [2, 4, 5, 8, 9, 17, 24, 31]. We show that super-exponential growth naturally emerges in a model of autocatalytic production of ribosomes, which determine the speed of cell elongation and surface area synthesis. This model allows us to derive the equations governing the dynamics of cell length and width, showing that super-exponential elongation in cell length occurs non-uniformly along the cell, while cell width fluctuates around a mean value that is dependent on the growth rate. Analysis of noise in growth and size parameters of *E. coli* cells [4] reveals strong intergenerational coupling of fluctuations in cell length and growth rate, whereas the fluctuations in cell width are independent of length. We then extend our dynamics into stochastic processes that accurately reflect the noise seen in experimental data as well as the correlations in model parameters across different growth conditions. In particular, we find that the dominant noise contributions come from model parameters that determine the rate of ribosomal synthesis, division protein synthesis, and cell surface area production. The resultant Langevin equations for cell length and width are capable of making predictions about the role of both intra-generational and inter-generational noise on the distribution of cell size and generation times in different nutrient conditions. Furthermore, using the single-cell level model we predict cellular strategies for ribosomal resource allocation and morphogenetic noise control in different growth conditions.

## II. RESULTS AND DISCUSSION

### A. Dynamics of cell growth

#### Single-cell data analysis

A common assumption in existing models of bacterial growth is that bacterial cells grow exponentially in size during the course of the cell cycle [2, 4, 5, 8, 9, 17, 24, 31]. We first re-examined this model by analyzing multi-genenerational growth and width data of single *E. coli* cells grown in the mother machine at steady-state under different nutrient conditions [4]. Parametrizing the geometry of rod-shaped *E. coli* cells by the pole-to-pole length *L* and width *w* (Fig. 1A-inset), we define the instantaneous growth rate as *κ*(*t*) = *L*^−1^d*L/*d*t*. To determine the overall trend in growth rate during the cell cycle, we averaged growth rate data across individual generations to obtain the average growth rate ⟨*κ*⟩ vs *t/τ*, where *t* is the time since birth and *τ* is the interdivision time. We found that the growth rate of the cell increases by ∼ 30% during the cell cycle in fast growth conditions (Fig. 1A), consistent with recent reports of increase in *E. coli* growth rate during the cell cycle [6, 32]. This trend in super-exponential growth, where the instantaneous growth rate *κ* increases over time, is preserved across different nutrient conditions (Fig. 1B), thereby invalidating existing models of purely exponential growth. As a test of whether this behavior is specific to *E. coli*, we applied the same analysis to rod-shaped *C. crescentus* cells grown in nutrient-rich media (PYE) at different temperatures [5, 11]. Our analysis confirms that the growth rate of *C. crescentus* cells increases during the cell cycle (Fig. 1C), suggesting that super-exponential growth is likely prevalent across different bacterial species.

**FIG. 1.**
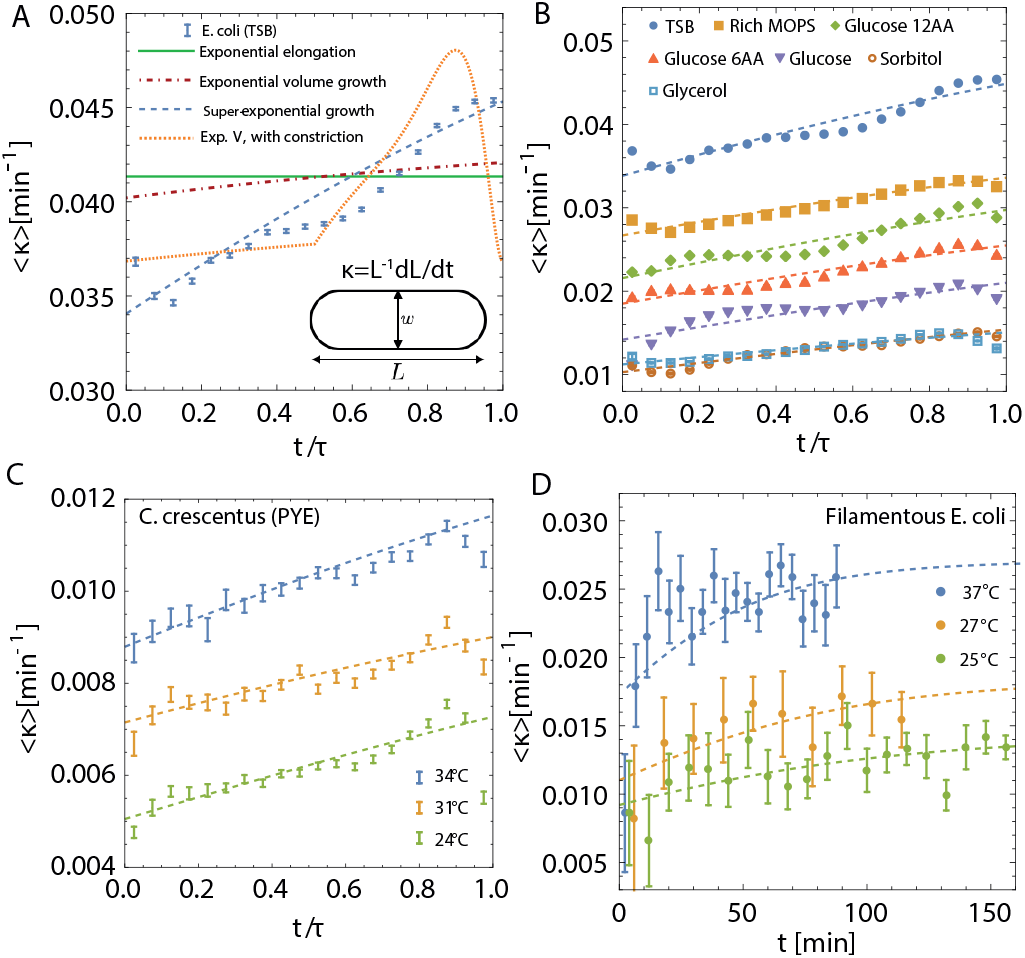
Super-exponential growth in *E. coli* and *C. crescentus* cells in different conditions. (A) Ensemble-averaged instantaneous growth rate of *E. coli* cells grown in TSB media at 37 *C* vs normalized time *t/τ*, where *τ* is cell cycle duration. Data taken from [4]. Error bars in all parts show ±1 standard error in the mean. The solid green line shows a fit of exponential length growth (1), dot-dashed red line represents a prediction from exponential volume growth (2), dashed blue curve shows a fit to the super-exponential growth model (3), and the dotted orange curve shows fit to exponential volume growth model with constriction dynamics. Fitting parameters: Model (1): *L*_0_ = 3.87 *μ*m, *κ* = 0.041 min^−1^, Model (2): *L* = 3.89 *μ*m, *k*_*V*_ = 0.044 min^−1^, Model (2) with constriction: *L*_0_ = 3.87 *μ*m, *k*_*V*_ = 0.04 min^−1^, Model (3): *L*_0_ = 3.99 *μ*m, *k* = 0.057 min^−1^, *λ* = 1.59 *μ*m. Cell width (*w* = 0.98 *μ*m) value is taken directly from experimental data. Inset: A simplified cell shape schematic for *E*. coli, defining the size parameters. (B) Fits of non-exponential growth model (3) to average growth rate data for seven different growth conditions grown at 37 *C*, taken from [4]. Error bars are negligible on the plotted scale. The values of *k* and *λ* for each condition are provided in Fig. 2B. (C) Fits of the super-exponential growth model (3) to average instantaneous growth rate vs *t/τ* of *C. crescentus* cells grown in PYE at three different temperatures. Data taken from [11]. (D) Time-dependent growth rate in filamentous *E. coli* cells grown in LB at different temperatures, presented in absolute time. Data taken from [6]. Dashed lines in (A-D) represent fits of the model in Eq. (3).

#### Testing data against existing models of cell growth

As discussed above, exponential elongation in cell length at a constant rate *κ*,

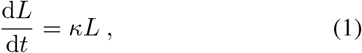

does not quantitatively capture cell cycle variations in growth rate despite its common use [2, 24]. Simplicity is the major upside of this model since microscopy data measure 1D and 2D geometrical quantities rather than cell volume. To understand the mechanistic origin of super-exponential growth, we first inquired if the increase in the rate of elongation in cell length is a geometric consequence of exponential growth in cell volume *V* [17, 33], d*V/*d*t* = *k*_*V*_ *V*, where *k*_*V*_ is the constant rate of volume growth. Assuming a spherocylinderical cell geometry (*V* = *πw*^3^/6 + (*L* − *w*)*πw*^2^/4), we derive the length equation

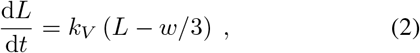

which can be interpreted as exponential growth of a portion *L* − *w*/3 of the cell’s total length. As seen in Fig. 1A, this model leads to a ∼ 3% increase in *κ* over time, which is an order of magnitude less than what is observed in the data. Thus, purely exponential growth in cell length or volume, defined by a one-parameter model, is not sufficient to capture the intra-generational dynamics presented in Fig. 1. Such simple models, however, are sufficient to describe phenomena on the inter-generational level such as cell size homeostasis and growth control [4, 9, 11].

It has recently been suggested that an increase in exponential growth rate could arise from the dynamics of cell constriction during septal growth [32]. We examined such a model by including the effects of constriction dynamics on the exponential growth of cell volume (see Fig S1 and Supplemetal Information for details). We found that constriction alone cannot explain the increase in growth rate of the cell throughout the cell cycle (Fig. 1A). In particular, the constriction model does not explain super-exponential growth until constriction initiation, exhibit a growth rate maximum prior to division, followed by a decrease in growth rate to the initial value as cell shape approaches that of two daughter cells. Furthermore, as we show later, super-exponential growth also occurs in filamentous cells that do not form a division septum.

#### Phenomenological model

Recent experiments have shown that the *E. coli* and *C. crescentus* cells do not grow uniformly along their length [11, 34], motivating a physical model where a portion *λ* of the total cell length does not grow:

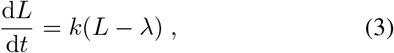

where *k* is the rate of cell elongation and the parameter *λ* can be determined by fitting the model to experimental data. This phenomenological model makes no assumptions about the spatial distribution of *λ*, but assumes that *λ* is time-independent. Attempts to model time-dependent *λ* results in inconsistencies with experimental data for *E. coli* or *C. crescentus*; increasing *λ* over time leads to sub-exponential growth, whereas a decrease in *λ* results in unbounded increase in growth rate that is inconsistent with data for filamentous cells (see below). Solving Eq. (3) for 0 ≤ *t* ≤ *τ* gives

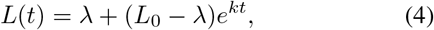

where *L*_0_ is the cell length at birth. Since *E. coli* behaves as an adder [4, 8, 9], cell division occurs when the cell length increments by a constant amount Δ: *L*(*τ*) = *L*_0_ + Δ. Thus the interdivision time is given by *τ* = *k*^−1^ ln (1 + Δ/(*L*_0_ − *λ*)). Upon division, each daughter cell is assigned a value of *λ* uncorrelated to that of the mother cell (Table II).

Comparison between the models given by Eq. (2) and (3) are provided in the data for cell length vs time (Fig S2A-B) and growth rate vs time (Fig. 1A-B). For the length data, goodness of fit tests reveal that model (3) most accurately captures the data (Fig S2C). With the model parameters determined from fitting the cell length data, model (3) is capable of fully capturing the increasing growth rate trend (Fig. 1A) across all nutrient conditions (Fig. 1B). Interestingly, we find that the average value of *λ* is larger than the average cell diameter in all growth conditions (Fig. 2B), suggesting that there are regions within the cylindrical portion that are non-growing in addition to the poles.

**FIG. 2.**
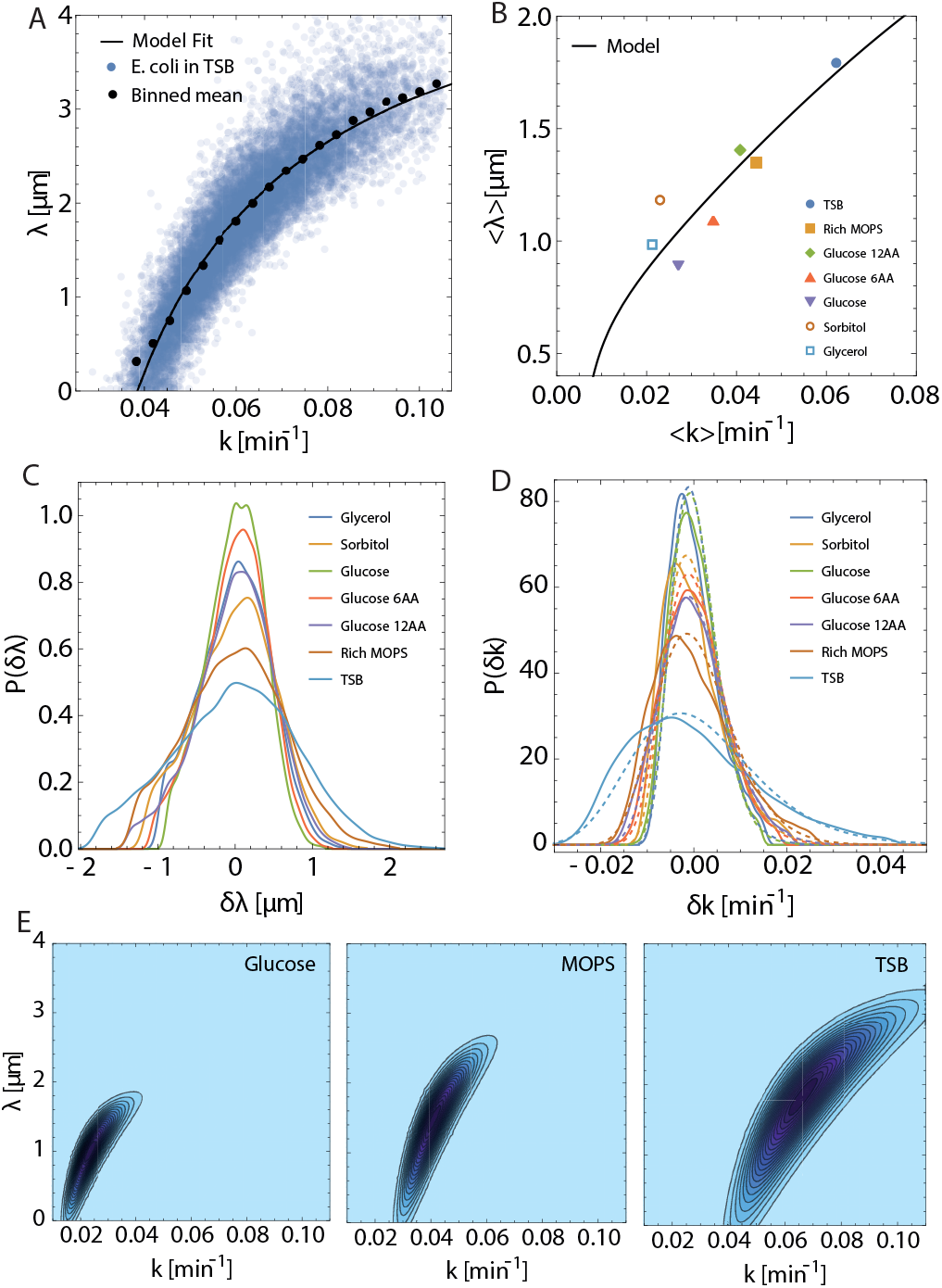
Intergenerational fluctuations and correlations in *E. coli* cell growth parameters. (A) Scatterplot showing the correlation between *k* and *λ* obtained by fitting super-exponential growth (9) model to *E. coli* cell length vs time data in TSB. The solid black curve represents a fit of Eq. (8) to the data. We remove outliers and the small number of generations in fast growth conditions with *λ <* 0 from further analysis (see Methods). (B) Ensemble-averaged ⟨*λ*⟩ vs ⟨*k*⟩ across different growth conditions. The black curve is a model prediction for ⟨*λ*⟩ as a function of ⟨*k*⟩ according to Eq. (9), with *L*_0_ = (1.50 *μm*) exp((16.19 min) *k*), *αR*_0_ = (0.84 *μm*) ⟨*k*⟩ exp((16.19 min) ⟨*k*⟩),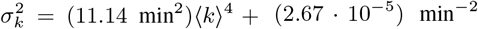. (C) Marginal probability distributions of *δλ* = *λ* − ⟨*λ*⟩ across different growth conditions. (D) Marginal probability distributions of *δk* = *k* − ⟨*k*⟩ for each growth condition shown as solid color curves. Dashed curves of the same color depict fits of log-normal distributions. Experimental data presented in (A-D) are taken from [4]. (E) Representative contour plots of the model predictions for the joint distribution *P* (*k, λ*), corresponding to mean growth rates in Glucose, MOPS, and TSB media. Darker blue indicates higher probability.

A key prediction of our model is that the growth rate of the cell saturates to a constant at longer times, such that cell growth becomes purely exponential. This is evident from the growth rate equation

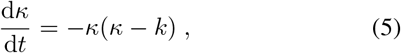

which predicts that *κ* ≈ *k* for *t* ≫ *κ*^−1^. To test this prediction, we analyzed the morphologies of filamentous *E. coli* cells [6] that have longer cell cycles due to impaired division. In agreement with our model, data show that the growth rate of filamentous cells increases initially and then saturates to a constant value over longer times (∼ 100 min at 25°C and 27°C), as shown in Fig. 1D. The timescale to reach exponential growth phase decreases with temperature, and thereby decreases with increasing growth rate, which is in agreement with our theory.

#### Mechanistic model of super-exponential growth

Our phenomenological model of super-exponential growth can be derived from an underlying molecular model that assumes cell length increases at a rate proportional to the abundance *R* of actively translating ribosomes

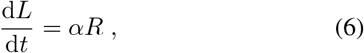

where *α* is the speed of cell elongation per ribosome, related to the translational capacity of the cell. This model is motivated by data that bacterial growth rate is linearly proportional to the mass fraction of actively translating ribosomes [35]. One could alternatively formulate Eq. (6) as the rate of volume growth proportional to ribosomes, resulting in re-scaling of the parameter *α* by geometric parameters of the cell. However, we choose to work with cell length as length data are directly measured in experiments whereas volume must be calculated using data for cell length, width, and geometric assumptions.

Since ribosomes are autocatalytic structures, the abundance of active ribosomes grows at a rate proportional to *R*

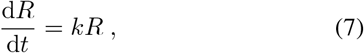

where *k* defines the rate of synthesis of ribosomal proteins. Solving for *L*(*t*), we arrive at the same equation as (4) with

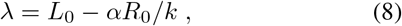

where *R*_0_ is the abundance of active ribosomes at cell birth, *k* is identified as the cell elongation rate (4), predicting that *λ* increases with *k*. In this form, *λ* is more clearly interpreted as the difference between cell’s actual length (*L*_0_) and the length of material synthesized by ribosomes (*αR*_0_*/k*) at birth. Thus, super-exponential (*λ >* 0) occurs when there is a mismatch between cell geometry and the initial protein synthesis capacity of the cell.

Deriving Eq. (3) from an underlying model of ribosome synthesis is important to its validity. Rather than introducing the two-parameter model in Eq. (3) as an experimentally motivated ansatz, we use a pre-existing understanding of ribosome synthesis to describe cell growth using two physiological parameters: translation speed *α* and ribosome synthesis rate *k*. These parameters in turn define the rate of cell growth and the portion of non-growing region of the cell, *λ* (Eq. 8).

### B. Intergenerational Fluctuations in Cellular Growth Parameters

While the deterministic growth model described by Eqs. (6)-(7) accurately describes the average growth dynamics of the cell, there are significant fluctuations in model parameters across different generations and growth conditions. Understanding these parameter variations is essential for predictive modeling of stochastic growth dynamics. To obtain a quantitative understanding of the noise in the cellular growth parameters between individual generations, we move from fitting ensemble-averaged data to individual generations within each growth condition for *E. coli* (Fig S2A) [4]. Fitting our effective growth model (3) to cell length data for each generation provides a pair of values for the elongation rate, *k*, and the effective length of the non-growing region, *λ*. From these fits, we find a positive correlation between *k* and *λ* within each growth condition, as shown by the scatterplot in Fig 2A. The mean trend in the correlation between *λ* and *k* is accurately captured by Eq. (8), which predicts that *λ* increases with *k*. Furthermore, we observe a strong positive correlation between the population means ⟨*λ*⟩ and ⟨*k*⟩ across nutrient conditions (Fig. 2B). To understand the origin of this correlation, we compute the relationship between ⟨*λ*⟩ and ⟨*k*⟩ using (8) under a small-fluctuation approximation *σ*_*k*_*/k* ≪ 1,

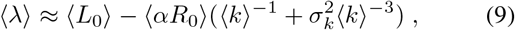

where *σ*_*k*_ is the standard deviation in *k*. In the above equation, both ⟨*L*_0_⟩ and ⟨*αR*_0_⟩ are functions of ⟨*k*⟩. Since average cell size increases exponentially with the nutrient-specific growth rate [14, 36], we assume an exponential form for the dependence of ⟨*L*_0_⟩ on *k* that we determine by fitting experimental data [4] (Fig S3A). Furthermore, since ribosome mass fraction increases linearly with growth rate [35], we fit ⟨*αR*_0_⟩ / ⟨*L*_0_⟩ to a linear function of ⟨*k*⟩ that captures the data very well (Fig S3B). With these fitted functions, we can model ⟨*λ*⟩ as a continuous function of ⟨*k*⟩ (Fig. 2B).

Next we turn to modeling the distributions of *λ* and *k* such that the intergenerational correlations are accurately captured. Marginalizing the joint distribution of *k* and *λ* obtained from experimental data, we find that the probability distributions of *δk* = *k* − ⟨*k*⟩ and *δλ* = *λ* − ⟨*λ*⟩ are skewed right and left, respectively, with increasing variance as ⟨*k*⟩ increases (Fig. 2C-D). We observe that the variations in *k* are reasonably well approximated by a log-normal distribution, as shown in Fig. 2D. This is not unexpected as the cellular elongation rate *k* is an accumulation of growth at many individual sites on the surface of the cell. However, the distribution of *λ* is slightly more difficult to model analytically (Fig. 2C). The unusual shape for *λ* distribution arises from the noise in *L*_0_, *k, α* and

*R*, through the relation defined in Eq. (8). Even under Gaussian noise approximations, an exact analytical distribution for *λ* does not exist. However, assuming small fluctuations about the mean values, we find that

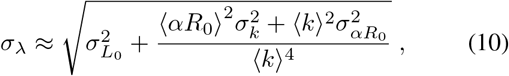

where *σ*_*i*_ is the standard deviation in parameter *i*. Interestingly, larger values of ⟨*αR*_0_⟩ leads to greater contribution of the noise in *k*, whereas larger *k* increases the contribution of the noise in *αR*_0_. An approximate analytical form the joint distribution *P* (*k, λ*) can be constructed by transforming the variables *k* and *λ* to remove the skew in the distribution: *K* = ln *k* and 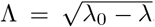, where *λ*_0_ is a chosen reference value. The transformed variables *K* and Λ are normally distributed, allowing us to construct a correlated multivariate normal distribution, ⟨*P*⟩ (*K*, Λ) (see Methods), whose parameters can be fitted with appropriate mathematical functions of the average elongation rate *k* (Fig S4). Representative contour plots of the joint distribution *P* are shown in Fig. 2E for three different growth conditions.

### C. Cell Width Maintenance During Growth

While cell length increases during the cell cycle, the width of rod-shaped *E. coli* fluctuates about a mean value *w*_0_ [4]. We consider two levels to these fluctuations: fluctuations of cell mean width *w*_0_ about the population mean ⟨*w*_0_⟩, and temporal fluctuations in cell width *w* about its mean *w*_0_ during any given generation. Fig. 3 characterizes the statistical properties of *w*_0_ as a function of the elongation rate *k*. Not surprisingly, ⟨*w*_0_⟩ increases exponentially with ⟨*k*⟩ at the same rate as cell length (Fig. 3A). This is a consequence of *E. coli* cells maintaining a constant aspect ratio (on average) in different nutrient conditions [19]. The distribution of *w*_0_ can be reasonably well approximated by a Gaussian, with the standard deviation of *w*_0_ increasing with ⟨*k*⟩ (Fig S5A). In addition, the coefficient of variation in *w*_0_ is roughly constant with changing growth conditions (Fig S5B). As a result, the probability distribution of *δw*_0_/ ⟨*w*_0_⟩, where *δw*_0_ = *w*_0_ − ⟨*w*_0_⟩, can be collapsed onto a single Gaussian distribution (Fig. 3B).

**FIG. 3.**
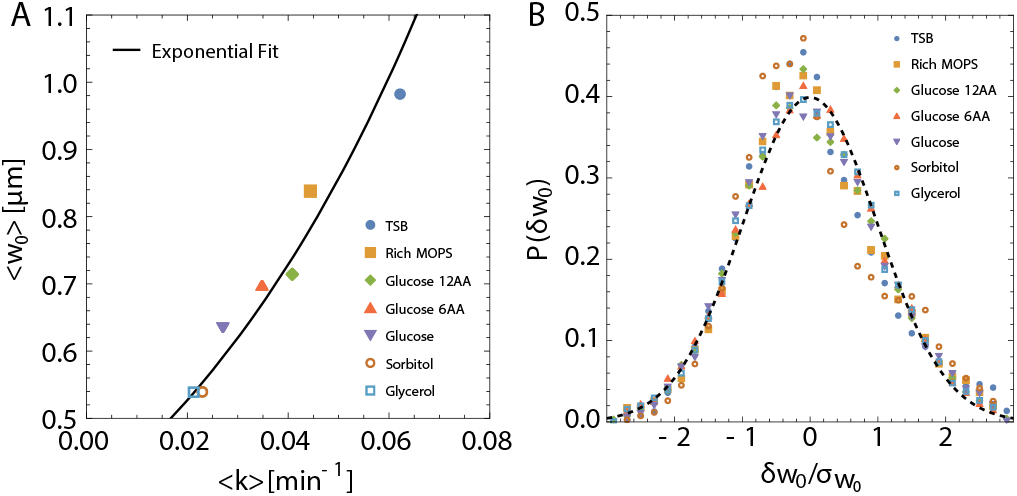
Intergenerational variations in *E. coli* cell width across growth conditions. (A) Average width ⟨*w*_0_⟩ vs ⟨*k*⟩ across growth conditions. The solid black line shows that ⟨*w*_0_⟩ increases exponentially with ⟨*k*⟩, keeping a fixed length-width aspect ratio (⟨*w*_0_⟩ = 0.25 ⟨*L*_0_⟩ = (0.38 *μm*) exp((16.19 min) ⟨*k*⟩)). (B) Probability distributions of the intergenerational fluctuations *δw*_0_ = *w*_0_ *w*_0_ across growth conditions scaled by their respective standard deviations 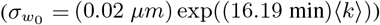. The dashed curve shows a universal Gaussian fit to the scaled data. Data are taken from [4].

To arrive at a general equation that describes the maintenance of cell width and its fluctuations around a mean value, we begin with the model that the surface area *S* of the cell is synthesized at a rate proportional to ribosomal abundance *R*,

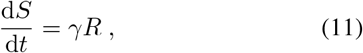

where *γ* is the rate of synthesis of surface material. This model stands in contrast to a recently proposed model of surface area synthesis in proportion to cell volume [17], but conceptually similar since surface material is assumed to be produced in the cytoplasm. Using a spherocylindrical geometry of the cell with pole-to-pole length *L* and width *w*, we have *S* = *πwL*. Using d*L/*d*t* = *αR* together with Eq. (11), we arrive at the width equation

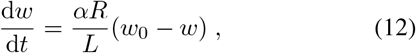

with *w*_0_ = *γ/πα*. Thus, *w* relaxes to the value *w*_0_ at steady-state, with *R/L* approaching the value *k/α*. Since the coefficient of variation in *w*_0_ is roughly constant across growth conditions (Fig 3B), it is reasonable to assume that the coefficients of variation in parameters *α* and *γ* are also maintained constant across different growth conditions.

Expressing *R* in terms of *L* and *λ* yields an alternative form for the width equation

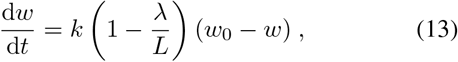

which showcases the asymptotic approach to d*w/*d*t* ≈ *k*(*w*_0_ − *w*), without considering the dynamics of *R* directly. This approach occurs on the timescale of the generation time *τ*, taking around 3*τ* to stabilize (Fig S5C).

### D. Intergenerational Dynamics and Correlations in Model Parameters

The dynamics of a bacterial cell in each generation are defined by Eq. (3) and (13), characterized by only three parameters, *k, λ*, and *w*_0_. These parameters vary across generations and growth conditions. To define the intergenerational dynamics, we expand our model to account for the rule of cell division. Cell growth is coupled to division such that the production rate of division proteins *X* (e.g., FtsZ) is proportional to the amount of active ribosomes [37, 38],

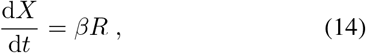

where *X* is the division protein abundance and *β* is the rate of synthesis of division proteins per ribosome. Cells divide once a threshold amount of division proteins, *X*_0_, are synthesized during the cell cycle [13, 22, 37, 38]. This leads to an adder model for cell size control, as relevant for *E. coli* cells [12], such that the added cell length (Δ) during each division cycle is constant and given by

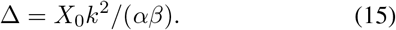

Next, we turn to computing the intergenerational correlations in model parameters, in order construct a stochastic model for cell width dynamics as it evolves through cycles of birth, growth and replication. There are two levels at which we must address the correlations in our model parameters: correlation between mother and daughter cells and population-wide correlation. In Table I, we list the intergenerational correlations in model parameters for each growth condition. As shown earlier in Fig. 2A, there is a strong positive correlation between *λ* and *k*. However, we do not observe a correlation between either *k* or *λ* with *w*_0_. This simplifies our model immensely since the length and width equations can now be simulated independently. It is interesting to note that there is a slight positive correlation between *λ* and the initial cell length *L*_0_, which decreases with increasing ⟨*k*⟩. This makes intuitive sense since *λ* is a portion of *L*_0_, and as *L*_0_ increases with growth rate the possible values of *λ* are less constrained. In other words, the contribution of *αR*_0_ in (8) is more pronounced for faster growing cells. We do not observe any substantial correlations between the added size Δ and the growth parameters *k* and *λ*.

**TABLE I.** Intergenerational correlations of model parameters

| Condition | $\rho(k, \lambda)$ | $\rho(\lambda, w_0)$ | $\rho(k, w_0)$ | $\rho(\lambda, L_0)$ |
| --- | --- | --- | --- | --- |
| Glycerol | 0.86 | -0.07 | -0.05 | 0.26 |
| Sorbitol | 0.84 | -0.09 | 0.04 | 0.20 |
| Glucose | 0.88 | -0.07 | 0.01 | 0.21 |
| Glucose 6AA | 0.89 | 0.02 | 0.12 | 0.21 |
| Glucose 12AA | 0.92 | -0.02 | 0.09 | 0.14 |
| Rich MOPS | 0.87 | -0.07 | 0.05 | 0.07 |
| TSB | 0.89 | -0.16 | -0.03 | 0.10 |

In Table II we examine the correlations in model parameters between successive generations. As expected from the adder model [12], there is a positive correlation in subsequent values of *L*_0_, with a correlation coefficient ≈ 0.5. Despite the slight positive correlation between *L*_0_ and *λ* at the population level (Table I), we see no correlation in *λ* values between successive generations. Furthermore, there is no correlation between subsequent generation values of *k*, so both *k* and *λ* can be drawn independently for each generation from the joint distribution *P* (*k, λ*) (Eq. 26). However, cell mean width is highly correlated between the mother and the daughter cell, such that the mean width in successive generations are related as

**TABLE II.** Correlations between successive generations

| Condition | $\rho_{n,n+1}(k)$ | $\rho_{n,n+1}(\lambda)$ | $\rho_{n,n+1}(L_0)$ | $\rho_{n,n+1}(w_0)$ |
| --- | --- | --- | --- | --- |
| Glycerol | 0.03 | 0.05 | 0.54 | 0.77 |
| Sorbitol | 0.00 | -0.01 | 0.56 | 0.59 |
| Glucose | 0.01 | 0.04 | 0.55 | 0.75 |
| Glucose 6AA | 0.00 | 0.03 | 0.51 | 0.81 |
| Glucose 12AA | 0.00 | 0.05 | 0.52 | 0.78 |
| Rich MOPS | 0.00 | 0.01 | 0.48 | 0.72 |
| TSB | 0.00 | 0.07 | 0.50 | 0.82 |

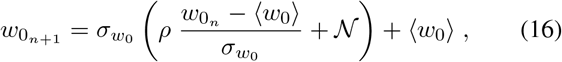

where *ρ* is the correlation between 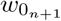 and 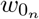, and 𝒩 is a Gaussian with mean 0 and variance 1 − *ρ*^2^. Further details about intergenerational mechanics are left to the Supplemental Methods. With all the intergenerational correlations and division mechanics accounted for, we now turn to developing the Langevin equations for cell length and width that can accurately capture intra-generational correlations and fluctuations in cell morphology and growth rates.

### E. Stochastic Length and Width Dynamics

To simulate the stochastic growth and size dynamics of single cells, we account for the noise in model parameters within each generation. We derive the Langevin equations governing the dynamics of cell length and width by considering noise in the parameters defining the deterministic equations (6)-(7), and (11). To this end, we write each parameter *q* (*α, k*, or *γ*) as 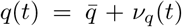, where 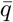 represents the average value of the parameter within a generation andν_*q*_(*t*) represents the time-dependent intra-generational fluctuations in *q*, assumed to be Gaussian white noises with zero mean. The assumption of white noise is motivated by the observation that the temporal fluctuations in cell length and width are uncorrelated in experimental data.

#### Width dynamic

– Considering stochastic variations in the parameters *α* and *γ*, we write the equation for cell width (12) as

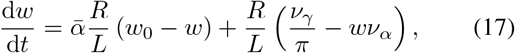

where 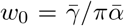. Considering the deviations from the cell mean width *δw* = *w* − *w*_0_, we find

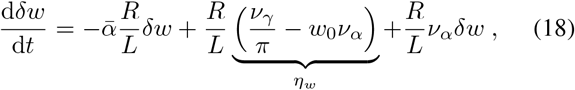

where *η*_*w*_ is a time-uncorrelated Gaussian additive noise with amplitude 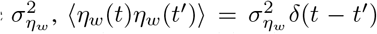. In addition, fluctuations in *α*, characterized by the noiseν_*α*_, contributes to multiplicative noise in the width equation. The interplay between additive and multiplicative noise with a restorative drift has been studied recently in the context of bacterial shape control [27]. Although our equation differs from [27], some of the conceptual results in [27] hold here as well. Crucially, the effect of increased multiplicative noise on the distribution of *δw* is to not only increase the spread but also to add a pos-itive skew to the distribution. In other words, neglecting the multiplicative noise (*ν*_*α*_ = 0) to write

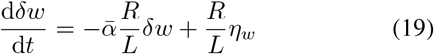

is a good approximation when *δw* is Gaussian. As a result, *δw* follows an Ornstein-Uhlenbeck process [39]. From the experimental data, we do not observe a skew to *δw* distribution and hence this approximation is justified. Neglecting the multiplicative noise is equivalent to neglecting intra-generational variations in *α* compared to *k* and *γ*.

Integrating Eq. (18) would require keeping track of the values of *R* as it fluctuates during growth. While this mathematically poses no difficulty, we do not currently have experimental data for the dynamics of *R*(*t*), so their values cannot be benchmarked in simulations. To circumvent this issue, we eliminate *R* to recast the width equation in terms of *L*:

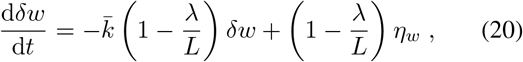

which can be integrated forward using information available from data. In the case of purely exponential growth in length (*λ* = 0), (20) takes the simple form

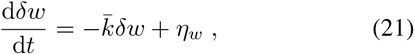

showing that width fluctuations decay over a timescale 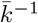 that is set by the elongation rate. Since (1 − *λ/L*) scales both the restoring force and the noise in the width equation, despite (1 − *λ/L*) increasing throughout the cell cycle we see no qualitative or quantitative differences between width trajectories generated by model (20) or (21) (Fig S 6). In other words, super-exponential elongation in length does not strongly affect stochastic width fluctuations within a generation.

#### Length dynamic

– The langevin equation for cell length can be derived using Eq. (6),

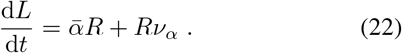

By eliminating *R* and neglecting the intra-generational noise in *α* (as justified earlier) we can recast the above equation in terms of ℓ= *L* − *λ*:

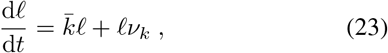

with multiplicative Gaussian white noiseν_*k*_, where ⟨*ν*_*k*_⟩ = 0, 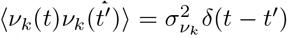, and 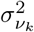 is the noise amplitude. The above equation may be recognized as the Black-Scholes model in stochastic form [39]. As for stochastic variations in *λ* within a generation, it follows from Eq. (8) that

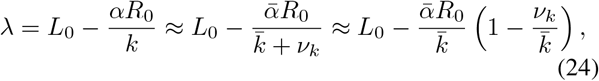

which is approximately constant in the small fluctuation limit 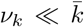. This is implicit in the assumption of writing the noise termν_*k*_ in Eq. (23). Rather than modeling *k* stochastically and updating *L* deterministically from *k*, we consider the fluctuations in *k* to be Gaussian around a mean 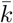, with no memory at each timestep during the cell cycle.

#### Ribosome dynamics

– While we do not directly model *R* to predict cell length and width dynamics, the stochastic dynamics of *R* during the cell cycle can be derived by considering noise in *k*,

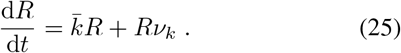

In other words, we predict a multiplicative noise in *R* with an amplitude 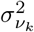 that can be determined from the measurements of length fluctuations. To model intergenerational dynamics, we note that an adder mechanism for cell length control implies an adder model for ribosome homeostasis, such that *R*_*n*_(*t* = *τ*_*n*_) = *R*_*n*_(*t* = 0) + *k*Δ_*n*_*/α*, where *n* is the generation index, *R*_*n*_ is the abundance of actively translating ribosomes, Δ_*n*_ is the added length and *τ*_*n*_ is interdivision time in generation *n*. By fitting cell length data, we can determine the parameters *α, k* and Δ to predict ribosome dynamics during the cell cycle (Fig S7A). The predicted dynamics show that following division, a certain amount of ribosomes is removed from the actively translating pool *R* for the daughter cell, such that 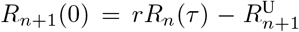, where *r* is the division ratio and 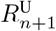 is the amount of ribosomes removed from the active ribosome pool in generation *n* + 1. This is a model prediction rather than an assumption, as *R*^U^ is determined from fitting data. We note that a non-zero *R*^U^ is necessary for super-exponential growth in consecutive generations, as observed in data (Fig S 2D), otherwise a cell would grow exponentially at a constant rate following division. There are several possible underlying explanations for non-zero *R*^U^, including fluctuations in free ribosome abundance and degradation, as described in the Supplemetal Information (Fig S7B).

#### Langevin simulations

With all the model parameters and their intra-generational and intergenerational fluctuations determined from experimental data, we can simulate the Langevin equations (20) and (23) in different growth conditions (see Methods). Fig. 4 shows representative trajectories of cell length and width resulting from such a simulation in fast and slow growth conditions. For both length and width, the simulations reproduce short timescale fluctuations within a given generation (4A and 4C), as seen in experimental data [4]. Longer timescale intra-generational length fluctuations about a smooth fit of (4) are also present, often spending ∼10-20% of the cell cycle above or below the average dynamics. The fluctuations about the intra-generational width mean are more pronounced than what we see for length dynamics, often spending more than ∼50% of a cell cycle without crossing the mean, consistent with experimental data [4]. At the intergenerational level, length fluctuations are regulated by the adder model (4B) but the fluctuations in width dynamics (4D) are worth noting. Based on our observed mother-daughter correlations in *w*_0_ (Table II), there is typically a large change in *w* during the division process compared to the intra-generational fluctuations.

**FIG. 4.**
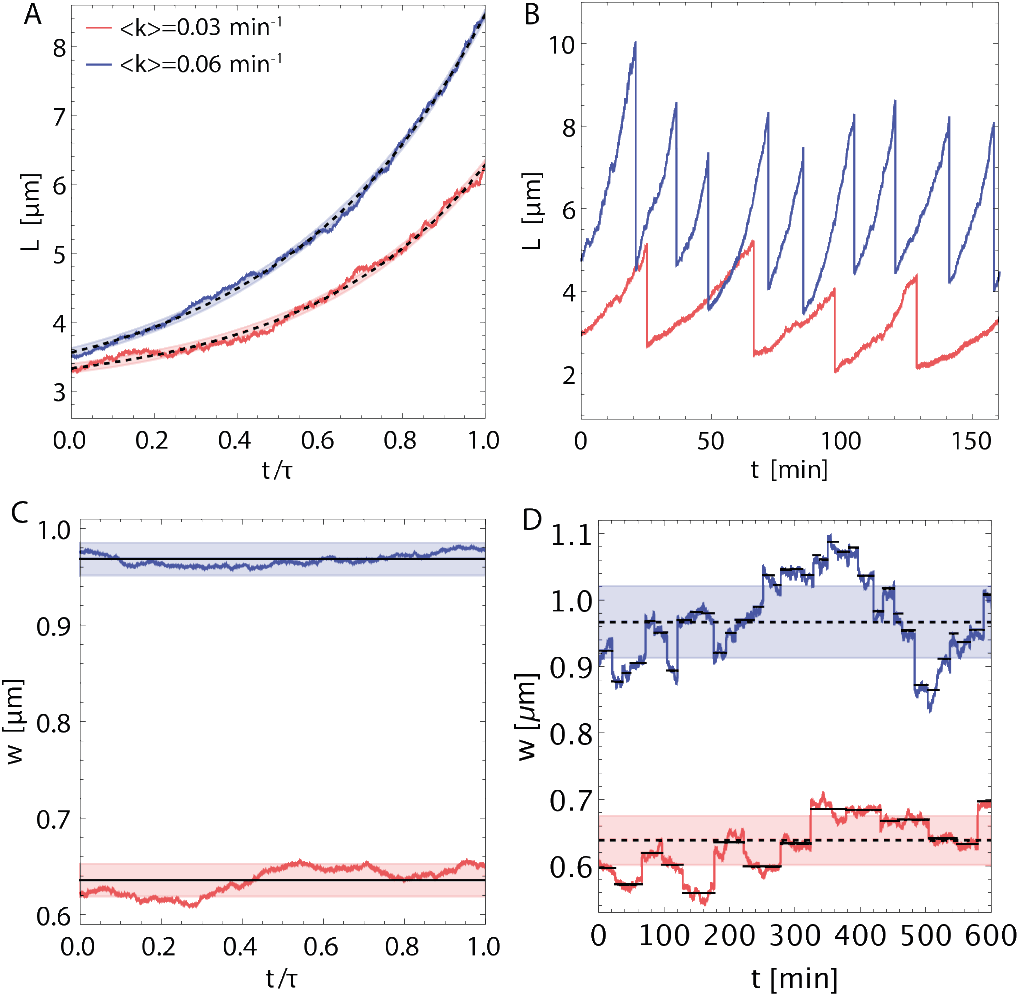
Langevin simulations for stochastic cell size dynamics in fast and slow growth conditions. For (A-D), blue trajectories correspond to a relatively fast growing condition with ⟨ *k* ⟩ = 0.06 min^−1^ and red to a relatively slow growing condition with ⟨*k*⟩ = 0.03 min^−1^. (A) Length vs normalized time *t/τ* for a single generation, where *τ* is cell cycle duration. The dashed black curve is a fit of deterministic super-exponential growth (3). The transparent bands surrounding each deterministic curve represent the standard deviation in length fluctuations (*σ*_*δL*_ = 0.066 *μm*). (B) Length vs absolute time for several generations. (C) Width vs *t/τ* for a single generation in normalized time. The solid black shows individual cell mean width. The transparent band around each cell mean width line represents the standard deviation in width fluctuations (*σ*_*δw*_ = 0.017 *μm*). (D) Width vs absolute time for several generations. Cell mean width is represented with solid black lines while population mean width is represented by dashed black. The transparent band around each population mean width line represents the standard deviation in intergenerational fluctuations 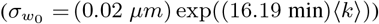.

The Langevin model can be used to generate predictions about the distribution of cell size and interdivision time. In Fig. 5, we simulate how the noise in cell cycle time (A) and initial cell size (B) changes as a function of the noise in growth at the intra- and intergenerational levels. Noise in added length Δ is propagated through from *k* according to Eq. (15). We observe that, while both intergenerational noise (*σ*_*k*_, *σ*_*λ*_) and intra-generational (*σ*_*νk*_) noise have an impact on the fluctuations in generation time, the effects of the intergenerational noise are more pronounced. Assuming symmetric division, cell size distribution at birth is controlled entirely by Δ and, while Δ does fluctuate in time withν_*k*_, the time-uncorrelated fluctuations in 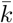 are too small to have an effect comparable to the cell-to-cell variation in Δ.

**FIG. 5.**
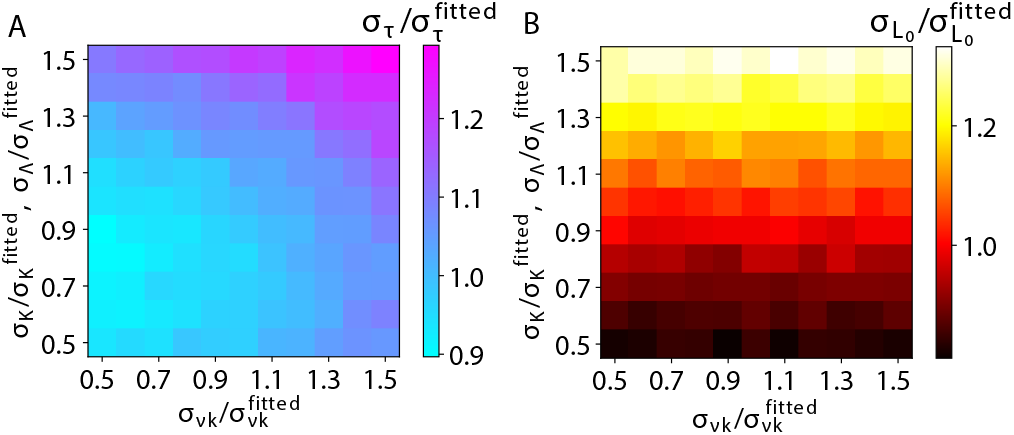
Noise in cell cycle time and cell size propagates from noise in growth parameters. (A) A colormap showing standard deviation in cell cycle time *τ* (*σ*_*τ*_) predicted by our stochastic simulations, as a function of intergenerational noise (*σ*_*K*_, *σ*_Λ_) and intra-generational noise (*σ*_*νk*_). Each axis is normalized by dividing the varied parameter(s) by the standard value(s) fitted to the data, and the scale of *σ*_*τ*_ is likewise normalized by the unperturbed value. Each parameter is varied ±50% along each axis. Bins sample 5000 generations for a single cell. (B) A colormap showing normalized standard deviation in initial cell size 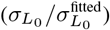, predicted by our simulations, as a function of intergenerational and intra-generational noise.

### F. Single-cell level resource allocation and noise control strategies

The kinetic model we developed for stochastic cell growth and size control can be used to derive single-cell level strategies for ribosomal resource allocation and morphogenetic noise control in different nutrient conditions. Previous studies on bacterial growth physiology at the population-level have revealed how bacteria allocate resources between ribosomal and metabolic protein synthesis in different nutrient conditions and under antibiotic treatments [35]. In particular, it was shown that there is a tradeoff between the mass fractions of ribosomal and metabolic proteins as nutrient conditions are varied [35]. Here we ask how such nutrient-dependent tradeoffs arise at the single-cell level between the allocation of cellular resources for growth, division and cell shape maintenance. Furthermore, we inquire how cells regulate noise in different physiological parameters as nutrient conditions are varied. Fig. 6A summarizes the main components of our model as defined by Eqs. (6)-(7), (14), and (11). Specifically, ribosomal proteins are involved in four major tasks: (1) production of ribosomes at a rate *k*, (2) cell elongation at a rate *α*, (3) division protein synthesis at a rate *β*, and (4) surface area synthesis at a rate *γ*. All these rates are controlled by the nutrient-specific growth rate. While we can determine *k* by directly fitting cell length data, we cannot directly extract the absolute values of the rates *α, β*, and *γ* from data. We therefore define normalized synthesis rates (same physical units as *k*): *α*^*i*^ = *αR*_0_*/L*_0_, *γ*^*i*^ = *γR*_0_*/S*_0_, *β*^*i*^ = *β/*(*R*_0_*X*_0_), which can be determined from fitting *E. coli* cell length and width data [4] considered in prior sections.

**FIG. 6.**
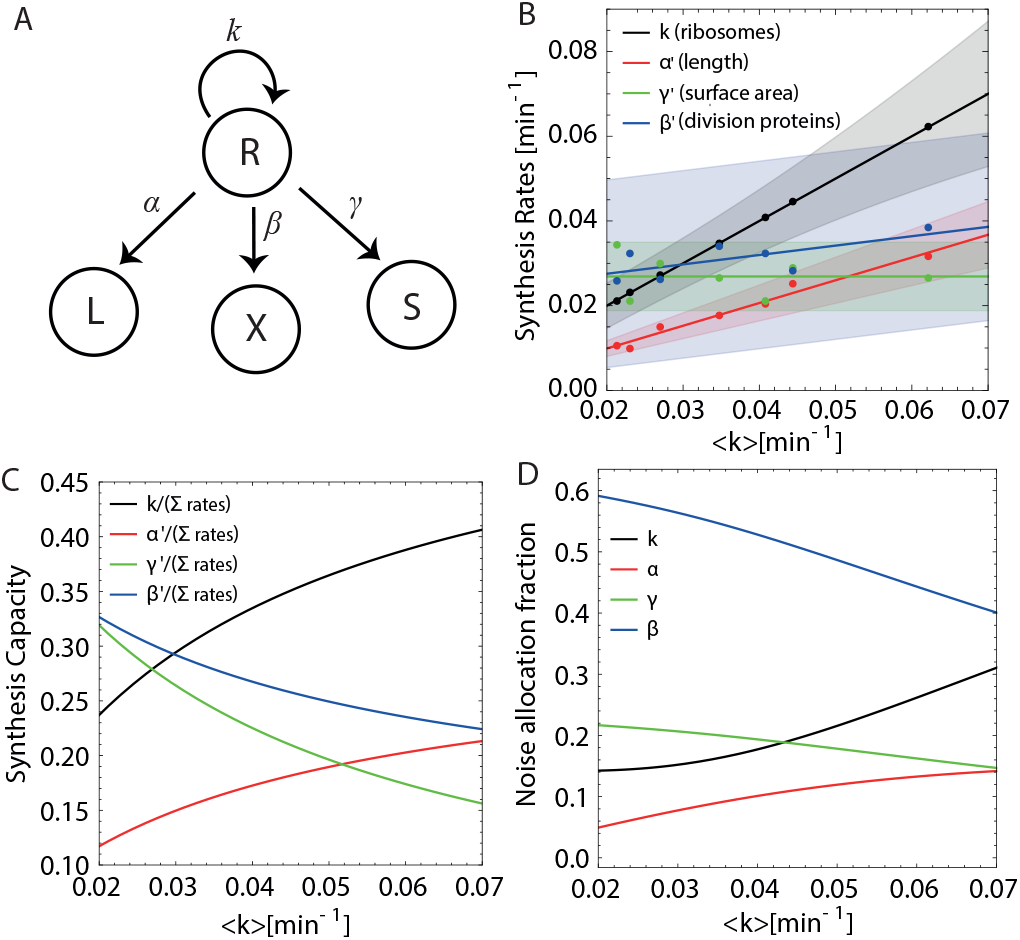
Cellular resource allocation and noise control strategies across nutrient conditions. (A) A network diagram for the underlying protein synthesis model. Ribosomes (*R*) responsible for synthesising new proteins do so in an auto-catalytic process (rate *k*) while also producing proteins necessary for growth and division (rates *α, β*, and *γ*). (B) Normalized synthesis rates (*α*^*l*^ = *αR*_0_*/L*_0_, *γ*^*l*^ = *γR*_0_*/S*_0_, *β*^*l*^ = *β/*(*R*_0_*X*_0_)) as a function of the mean elongation rate ⟨*k*⟩. Solid lines depict the mean values of the normalized rates, determined from fitting our model to experimental data (solid circles, showing mean values), and the transparent bands show one standard deviation of the corresponding distributions (see Methods). (C) Mean rates (as depicted in (B)) normalized by the sum of all rates for a given ⟨*k*⟩. (D) The noise (standard deviation) in each rate corresponding to the transparent bands in (B) normalized by the sum of all rates at a given ⟨*k*⟩.

From fits of our model to experimental data, we find that both ⟨*α*^*i*^⟩ and ⟨*β*^*i*^⟩ increase with ⟨*k*⟩ while ⟨*γ*^*i*^⟩ remains approximately constant (Fig. 6B). At the resolution of the data and our fitting, we see no change in the amplitude of noise in *γ*^*i*^ and *β*^*i*^, while the noise in *α*^*i*^ and *k* increases with ⟨*k*⟩. The increase in the rate of production of length material (⟨*α*^*i*^⟩) and the rate of division protein synthesis (⟨*β*^*i*^⟩) with ⟨*k*⟩ is consistent with data that both cell size and the rate of cell division increase with growth rates [13, 14, 37]. Since the rate of production of surface material remains approximately constant with increasing growth rate and cell size, it implies that surface-to-volume ratio decreases with growth rate, as seen in experimental data [14, 17, 19].

Given that most of the synthesis rates increase with ⟨*k*⟩, it is more insightful to interpret these results in the context of allocation of ribosomal resources to each these physiological tasks. We therefore introduce the rate allocation fractions, *ϕ*_*q*_ (*q* ∈ {*k, α*^*i*^, *β*^*i*^, *γ*^*i*^}), defined as the mean value of the rates ⟨*q*⟩, normalized by the sum of all mean rates (∑ ⟨*q* ⟩ = ⟨*k* + *α*^*i*^ + *β*^*i*^ + *γ*^*i*^ ⟩). As shown in Fig. 6C, both *ϕ*_*k*_ and *ϕ*_*α*_′ increases as ⟨*k* ⟩ increases, while *ϕ*_*β*_′ and *ϕ*_*γ*_′ decreases with *k*. It thus becomes clear that as nutrient-specific growth rates increase, cells allocate more ribosomal resources to producing more ribosomes and increasing cell size (length), while proportionally less resources are allocated to the production of cell division proteins and surface area synthesis. This trend can also be seen in noise allocation fractions determined from experimental data (Fig. 6D), where we find that the relative noise in ⟨*k* ⟩ and *α*^*i*^ increases while those of *γ*^*i*^ and *β*^*i*^ decreases with increasing *k*. In other words, more of the noise is present in the rate constants with greater allocation fraction. Taken together, these data show that there is a nutrient-dependent tradeoff between cellular resources allocated ribosome synthesis and cell size, and those that are allocated to synthesizing cell surface area and division proteins. These tradeoffs underlie the control of bacterial cell growth and morphology, and the regulation of noise in cellular growth and morphogenetic parameters.

## III. CONCLUSIONS

The predominant assumption for most of the history of bacterial growth modeling has been that exponential growth occurs at both the population and individual scales [12]. Our observations contradict this standard; we find super-exponential growth in cell size across a variety of nutrient and temperature conditions. We propose mechanistic models that account for the increasing exponential growth rate during the cell cycle – first in terms a phenomenological model of non-uniform cell envelope growth, and second in terms of a mechanistic model of autocatalytic ribosome synthesis. In the phenomenological model of non-uniform growth, the non-growing portion *λ* of the bacterial cell length can be interpreted using the mechanistic model as the mismatch between the cell’s geometry and initial capacity to synthesize proteins necessary for growth. Live-cell imaging of cell envelope growth pattern in *E. coli* during cell cycle progression would be necessary to test our interpretation of the non-uniform growth model. While superexponential elongation could also be captured by modeling the effects of constriction on exponentially growing volume of the cell, we find that the qualitative features of the constriction model are inconsistent with experimental data. Experiments quantifying septal growth dynamics in constricting *E. coli* cells is needed to directly test our model for constriction dynamics and study the effects division septum geometry on cell elongation.

Our model for cell growth dynamics comes with several parameters that have been calibrated based on experimental data. All of the parameter distributions that make up the dynamic models for cell growth, division, and width maintenance have been parametrized as functions of cellular elongation rate alone, allowing us to make quantitative predictions for how the average values and the noise in cellular parameters evolve in arbitrary growth conditions. Combining intergenerational noise factors with intra-generational stochastic differential equations, we uncover the degrees to which various underlying noise sources in growth and division processes contribute to overall noise in bacterial growth and morphogenesis.

While our stochastic growth model is developed primarily using single-cell data for *E. coli*, the modeling approach could be extended to other bacterial species with different morphological features and growth laws. In this context, it is pertinent to ask whether super-exponential growth is prevalent in other bacterial organisms, and not just limited to Gram-negative *E. coli* and *C. crescentus* cells. Analysis of single-cell growth data of gram-positive *Bacillus subtilis* cells [40] reveals a non-monotonic trend in growth rate during the course of the cell cycle (Fig S8A-B), as recently reported [33]. In particular, *B. subtilis* cells show a period of decelerated growth followed by a period of accelerated growth, irrespective of growth conditions. Our model is capable of capturing these behaviors with a time-dependent *λ* in Eq. (3), arising from time-dependent ribosome allocation, where *λ* increases during the first phase of decelerated growth and decreases during the second phase of accelerated growth (Fig S8C). These observations raise questions on how time-dependent changes in growth parameters are connected to cell cycle-dependent changes in cell envelope growth pattern and protein synthesis, for which experimental data are currently lacking. This time dependence has the potential to better capture the slight non-monotonicity observed in Fig. 1. A better understanding of how cellular growth parameters fluctuate during the course of the cell cycle would allow us to further refine our model assumptions and test theoretical predictions.

The mechanistic model based on ribosome synthesis and resource allocation could be directly tested in experiments measuring time-dependent bacterial proteome and ribosome synthesis during the cell cycle. While the production of ribosomes, cell division proteins and cell envelope proteins are accounted for in our simplified model, we neglect many metabolic proteins, transporters, and those that fall into the housekeeping/maintenance sector of the proteome. Future work taking into account these details could provide insights into the biochemical processes that are difficult to measure directly in experiments. The amount of ribosomes unused for growth, *R*^U^, which is required for super-exponential growth, is an interesting prediction of our model that can be tested in future experiments. Non-zero *R*^U^ could potentially result from ribosome degradation [41] or a temporary increase in free ribosome abundance following division, although the exact dynamics of these processes are not verifiable with current experimental data. Future experimental work measuring translation kinetics through single-cell ribosomal profiling and RNA sequencing as a function of cell cycle time would directly challenge or support this prediction. These experiments would help test our predictions that in nutrient-rich growth environments, cells allocate proportionally more of their ribosomes to cell elongation and ribosome production rather than the synthesis of surface area material and division proteins (Fig. 6C). While these predictions are derived at steady-state growth conditions, future work predicting intragenerational dynamics of cellular resource allocation, particularly in changing nutrient conditions, would be of particular interest to the growing field of single-cell physiology.

## METHODS

### Growth rate calculation

The *E. coli* strain for the data considered from [1] is K12 NCM3722 (not-motile derivative SJ202). The K-12 MC4100 strain is used for the filamentous *E. coli* cells [6]. All growth rate data analyzed in this paper is calculated from cited timeseries length data. We perform this calculation using a midpoint derivative approximation. While this calculation loses the first and last data points, it is less noisy than a left/right approximation. We have considered a running exponential fit (∼3 points) to extract *κ* but this method is also more noisy than the midpoint approximation. For population average data, as shown Fig. 1, we average in normalized time. The growth rate is approximated at the individual cell level and then each data point is placed into population bins according to where the data point falls in *t/τ* (the cell cycle time *τ* is specific to each cell cycle considered). We choose the number of bins in each case to be less than the average number of data points collected per cell cycle to avoid misrepresentation. Averages and error bars presented are then calculated for each bin with data from all cells. When considering individual generations rather than averages (section II B onward), fits of (4) are performed on directly the length data rather considering the processed growth rates.

When obtaining values of *k* and *λ* through fitting Eq. (4) to individual cell cycles, in addition to removing outliers with non-physical parameters, we neglect the fits that result in *λ <* 0. These results only occur in fast growth conditions such as TSB and MOPS, accounting for less that 5% of the total cell generations. While the interpretation of *λ <* 0 is non-physical in the model where *λ* is interpreted as the length of the non-growing region of the cell envelope (Eq. 3), *λ <* 0 is physically permissible in the context of the ribosome model (Eq. 8), implying sub-exponential growth. Since sub-exponential growth is not observed in experimental data, the negative values of *λ* result from fitting errors for data with high intra-generational noise.

### Distribution of model parameters

Constructing an analytical form for the joint probability distribution of growth parameters *λ* and *k* is challenging due to the skew in the distribution. To circumvent this difficulty, a commonly used approach is to transform the data to first remove the skew in the distribution. Given the log-normal nature of *k* distribution, ln *k* removes most of the skew from *k* while reflecting *λ* (around a chosen reference value *λ*_0_) followed by a square root is sufficient to remove the skew. Most functions commonly used to reduce skewness are defined only on a positive domain and are also more effective if used on distributions that start without a large offset from 0, which means we must be careful about the point around which we reflect *λ*. For a single dataset, an easy choice is to perform a reflection such that the maximum value becomes the minimum and vice versa. However, since we aim to fit this distribution as a function of *k*, choosing to reflect in this manner introduces a new parameter *λ*_0_ that can be fit from the data. It is worth noting that *λ*_0_ only affects the efficacy with which the data are normalized rather than a modification to the data themselves. We thus define the trans-formed variables *K* = ln *k* and 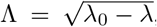, which are normally distributed, allowing us to construct the joint distribution

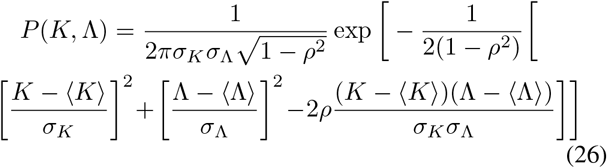

where the distribution parameters vary with growth conditions paramterized by ⟨*k* ⟩. As shown in Fig S 4, fitting appropriate mathematical functions for ⟨*K* ⟩, ⟨Λ ⟩, *σ*_*k*_, *σ*_Λ_, *λ*_0_, and the correlation function *ρ*(*K*, Λ), permits us to construct a correlated multivariate normal distribution for a given growth condition, determined by the parameter ⟨*k* ⟩. Representative contour plots of the joint distribution *P* are shown in Fig. 2E for three different growth conditions.

It is possible to expand this framework to include the slight positive correlation between *λ* and *L*_0_, as given in Table I. Considering a joint distribution including *L*_0_ in addition to *K* and Λ, we have

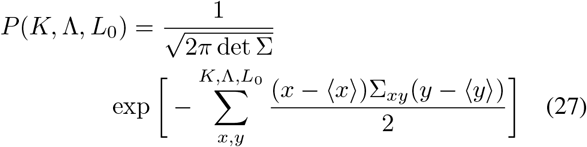

where σ denotes the covariance matrix. In the context of determining *K* and Λ for a new cell, (27) takes the fixed argument of the cell’s *L*_0_ determined by the division mechanism.

For parameters defining the ribosome synthesis model, their dependencies on mean elongation rates are given by: 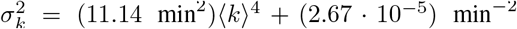, ⟨*α*^*i*^ ⟩ = 0.537*(k)* − 0.001 min^−1^, *σ*_*α*_′ = 0.120 ⟨*k* ⟩ − 0.001 min^−1^, ⟨*γ*^*i*^ ⟩ = 0.027 min^−1^, *σ*_*γ*_′ = 0.008 min^−1^, ⟨*β*^*i*^ ⟩ = 0.222 ⟨*k* ⟩ + 0.023 min^−1^, *σ*_*β*_′ = 0.022 min^−1^.

### Langevin model simulations

We briefly summarize the components that make up the Langevin model simulations. As an input, a function for mean elongation rate ⟨*k*(*t*) ⟩ is necessary. This fixes all distributions, as seen in Figs. 2, 3, S1-S4. For each generation, growth parameters *k* and *λ* are chosen according to our correlated joint distribution (27) in addition to a value of *w*_0_. We then integrate the stochastic differential equations (20) and (23) until division occurs, as prescribed by the adder model and a chosen Δ from fitted distribution to experimental data. We use a straightforward Euler-Maruyama method for the integration [42] using Itô calculus, neglecting the integration discretization intricacies that arise in the derivations of equations with non-constant diffusion. In each equation, the noise terms can be written as *η*(*t*) = *σξ*(*t*), where 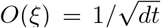. To perform the integration, we take *ξ*(*t*) = *dW* (*t*)*/dt* where *dW* is a zero-mean Gaussian with variance *dt* at each timestep (Weiner process). The noise amplitudes are determined by minimizing the difference between experimental and simulated width and length fluctuations, *σ*_*δw*_ (≈ 0.017 *μm*) and *σ*_*L*_, respectively (≈ 0.066 *μm*). We find that 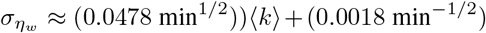 and 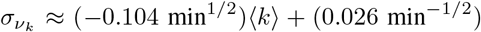. During cell division, length is split according to a Gaussian division ratio *r* and a new *w*_0_ for the daughter cell is chosen correlated to the old *w*_0_ according to (16). New values of *k* and *λ* are chosen uncorrelated to the previous generation and the cell cycle repeats. A sample output of the simulation is shown in Fig. 4 depicting results for slow and fast growing media.

## DATA AND CODE AVAILABILITY

Custom simulation code and analyzed data are available at https://github.com/BanerjeeLab/bacterial_growth_model.

## AUTHOR CONTRIBUTIONS

C.C and S.B. designed and developed the study. C.C. carried out simulations and analyzed the data. C.C. and S.B. wrote the article.

## ACKNOWLEDGEMENTS

We gratefully acknowledge support from the National Institutes of Health (NIH R35 GM143042) and the Shurl and Kay Curci Foundation. We thank Suckjoon Jun Lab (UCSD) for providing single-cell growth and shape data for *E. coli* and *B. subtilis* cells, Aaron Dinner and Norbert Scherer (University of Chicago) for providing data for *C. crescentus* cells.

