## Supplementary Information for "Super-exponential growth and stochastic size dynamics in rod-like bacteria"

### SUPPLEMENTAL METHODS

#### Modeling the effects of cell constriction on growth dynamics

Here we discuss the effects of constriction dynamics on the growth rate of a dividing cell. We begin with the equation for exponential volume growth: $\mathrm{d}V/\mathrm{d}t = k_V V$, where $k_V$ is the volumetric growth rate. Using a spherocylindrical geometry for the cell, with the length of the septal region along the cell's long axis given by $d$ (Fig S8A inset), the volume of the septal region is given by $V_s = (3w-d)d^2\pi/12$. Thus $V(L,w,d) = \pi w^3/6 + (L-w-d)\pi w^2/4 + (3w-d)d^2\pi/12$. Rearranging for $L$, we find that

$$L(t) = (L_0 - w/3)e^{k_V t} + w/3 + d(t) - \frac{d(t)^2}{w} + \frac{d(t)^3}{3w^2}. \tag{1}$$

Thus, given a mathematical form for $d(t)$, we can fit $L(t)$ and $\kappa(t) = (1/L)(\mathrm{d}L/\mathrm{d}t)$ to experimental data. Prior works have found that constriction does not start until around (or after) $t/\tau = 0.5$ [32]. With the initiation of constriction occurring at $t_c > 0.5$, we model $d(t)$ as a smooth function with $d(t < t_c) = 0$ and a fitting polynomial for $t > t_c$, subject to the boundary condition $d(\tau) = w$, i.e., the septum is fully formed at division (Fig S1A). The resulting fit is shown in Fig. 1A, showing the onset of super-exponential during the constriction phase as opposed to the entire cell cycle as seen in data. Furthermore, exponential volume growth with constriction requires the growth rate to fall back to the initial value during the same cell cycle, which does not match with each *E. coli* dataset addressed in this work (Fig S1C). Fig. S1B shows how $\mathrm{d}L/\mathrm{d}t$ changes with time during constriction, causing this non-monotonicity as the cell shape approaches that of two daughter cells.

#### Intergenerational Modeling of Bacterial Growth

Here we detail how the underlying variables of our growth model, ribosome abundance $R$, cell length $L$, surface area $S$, and division protein copy number $X$, change from one generation to the next and how they are affected by cell division. Following the adder model for cell size regulation, cell length at division is related to the cell length at birth as: $L_n(\tau) = L_n(0) + \Delta_n$, where $n$ is the generation index. Thus, assuming symmetric division, we obtain a recursion relation connecting cell length at birth at successive generations: $L_{n+1}(0) = (L_n(0) + \Delta_n)/2$. This relation ensures a very strong correlation between the birth sizes of subsequent generations such that the birth size approaches the mean added size. The adder model is equivalent to assuming that division occurs once a threshold $X_0$ of division proteins is reached, leading to the relation $\Delta_n = X_0 k_n^2/(\alpha_n \beta_n)$. For the purposes of our modeling and without loss of generality, we assume that the division proteins are used up during division such that $X_0(0) = 0$, which is also equivalent to assuming that $X$ is the amount of division proteins accumulated since birth. Given that we neglect shape change due to septum formation in our model, the way $S$ is handled during division is simply a consequence of $L_{n+1}(0)$, i.e. $S_{n+1}(0) = \pi w_{n+1}(0) L_{n+1}(0) = \pi w_{n+1}(0)(L_n(0) + \Delta_n)/2$, with the steady-state value $S(0) = \pi w_0 \Delta$.

An adder mechanism for cell length control implies an adder model for ribosome homeostasis, such that $R_n(\tau) = R_n(0) + k_n \Delta_n / \alpha_n$. Following symmetric division, $R_{n+1}(0) = \frac{1}{2} R_n(\tau) - R^{\mathrm{U}}_{n+1}$, where $R^{\mathrm{U}}_{n+1}$ is the amount of 'unused' ribosomes that are removed from the active ribosome pool $R$ following the division event. There are few potential physical explanations for a non-zero value of $R^{\mathrm{U}}$. It is possible that the rate of ribosome degradation peaks around the division event rather than remaining a time-independent quantity. Currently, experimental data to confirm or deny this is not available and such future work would be an interesting test of the model. Another possibility is that there is a temporary increase in free ribosome abundance, $R_{\mathrm{f}}$, following division. Consider a simple model of ribosomes transitioning between active ($R$) and inactive ($R_{\mathrm{f}}$) forms: $\mathrm{d}R/\mathrm{d}t = (k - d - k_-)R + k_+ R_{\mathrm{f}}$, $\mathrm{d}R_{\mathrm{f}}/\mathrm{d}t = k_- R + (k_+ - d)R_{\mathrm{f}}$, where $d$ denotes the passive degradation rate, $k_+$ and $k_-$ are he binding and unbinding rates of ribosomes to and from mRNAs. In this framework, a non-zero $R^{\mathrm{U}}$ could result from a temporary increase in $R_{\mathrm{f}}$, potentially due to a decrease in transcription rates and/or an increase in the rate that mRNA unbinds from ribosomes, following division. $R_{\mathrm{f}}$ then decays back into $R$ throughout the following cell cycle. Fig S7B illustrates these dynamics with parameters fit to data, as a refinement of the simple model of $\mathrm{d}R/\mathrm{d}t = kR$. Making physical assertions about these fitting parameters beyond a proof of concept is beyond the capacity of what can be inferred from cell length and width data.

Returning to the dynamics of $R$, we can write a recursion relation connecting ribosome abundance in successive generations: $R_{n+1}(0) = (R_n(0) + k_n \Delta_n / \alpha_n - 2R^{\mathrm{U}}_{n+1})/2$. Recalling $\lambda_n = L_n(0) - \alpha_n R_n(0)/k_n$, in terms of average dynamics (no fluctuations in $k$ and $\alpha$) we find that $\lambda_{n+1} - \lambda_n \approx \alpha R^{\mathrm{U}}_{n+1}/k - \lambda_n/2$. This expression indicates that a non-zero $R^{\mathrm{U}}$ is necessary for a non-zero $\lambda$ to persist. In terms of steady-state values, $L(0) = \Delta$, $R(0) = k\Delta/\alpha - 2R^{\mathrm{U}}$, and $\lambda = 2\alpha R^{\mathrm{U}}/k$. Fig S7 illustrates these values in experimental context for a single growth condition.

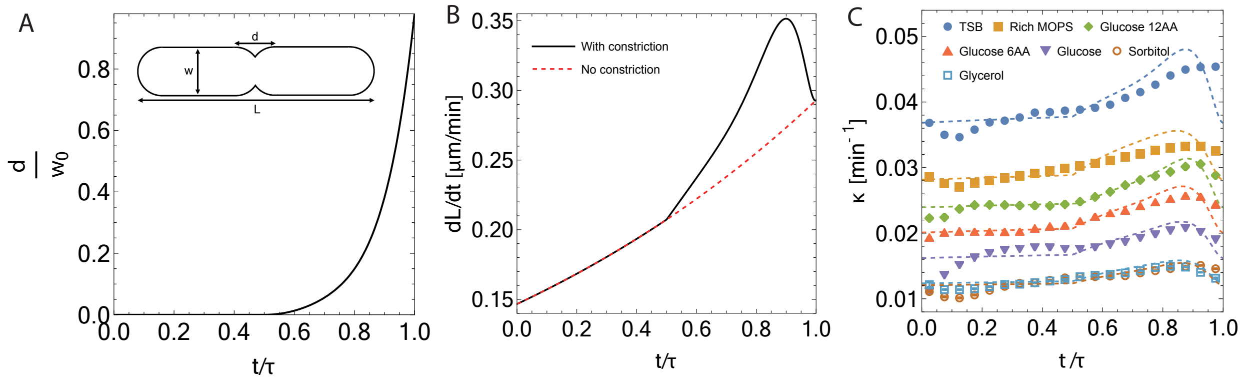


**Figure S** 1. **Effects of constriction on *E. Coli* growth dynamics.** (A) Length of septum $d$ (normalized by average cell width of the cylindrical portion), as a function of time. $d(t)$ is an 8th order fitting polynomial and constrained as described in the supporting methods. Inset: Schematic of a constricting cell. (B) The average rate of change in $L$ for cells grown in TSB at 37 $C$ taken from [4]. A fit of exponential volume growth is shown in black and the constriction volume growth model is shown in dashed red. (C) Fits of the constriction volume growth model (1) to average growth rate data for seven different growth conditions grown at 37 $C$, taken from [4]. Error bars are negligible on the plotted scale.

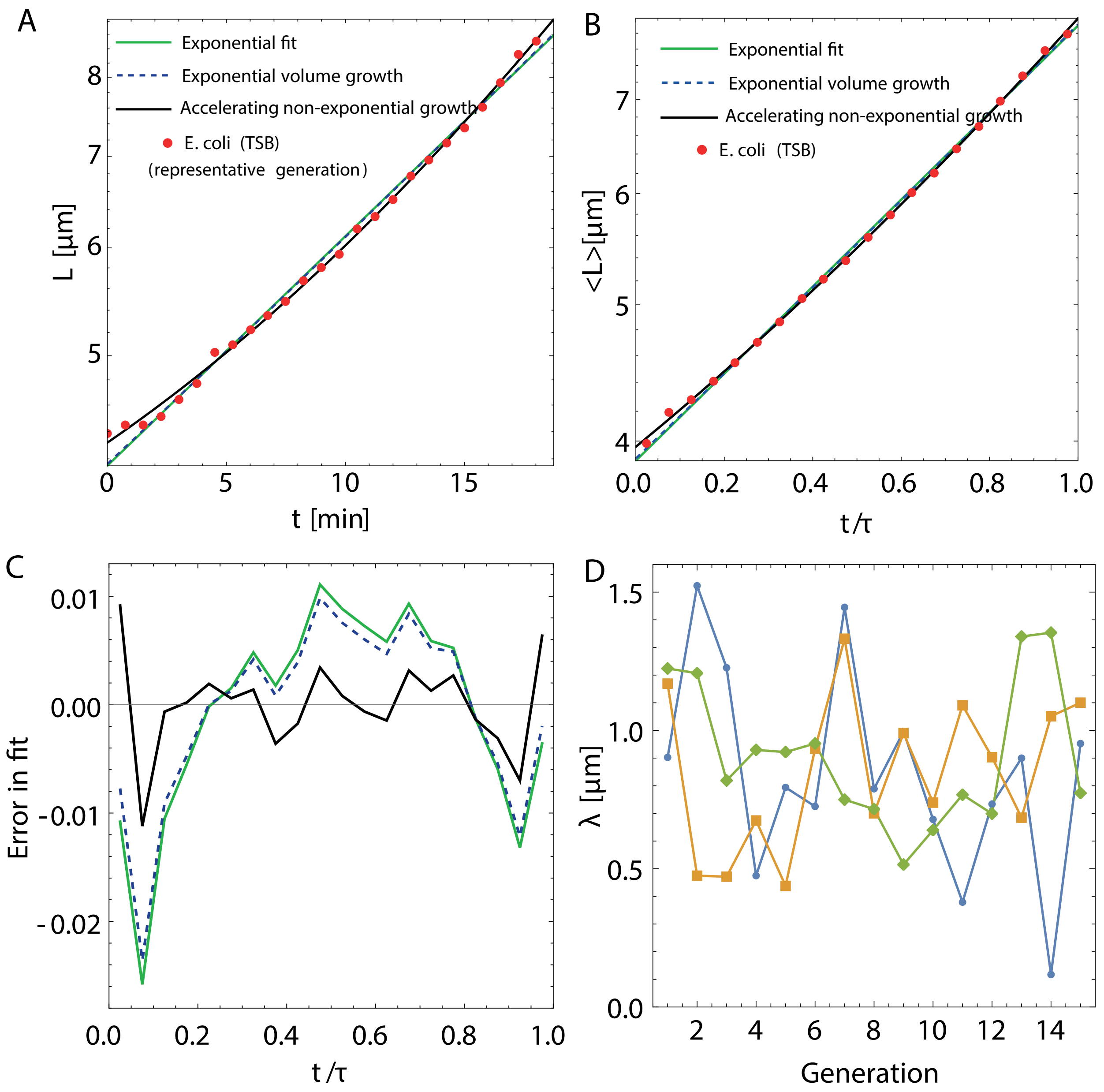


**Figure S 2. Cell length dynamics of *E. Coli*.** (A) Length of a representative cell cycle grown in TSB media at 37°$C$ vs absolute time fit with exponential growth in length (green), volume (dashed blue), and super-exponential growth rate model (black). (B) Ensemble-averaged length of *E. coli* cells grown in TSB media at 37°$C$ vs normalized time $t/\tau$, where $\tau$ is cell cycle duration. The average data is fit with exponential growth in length (green), volume (dashed blue), and super-exponential growth (black) with neglidgible error bars. (C) Error from each of the fits in (B) to the experimental data. (D) Evolution of growth parameter $\lambda$ across generations for three representative cells, showing that $\lambda$ is uncorrelated between successive generations. Data for (A-D) taken from [4].

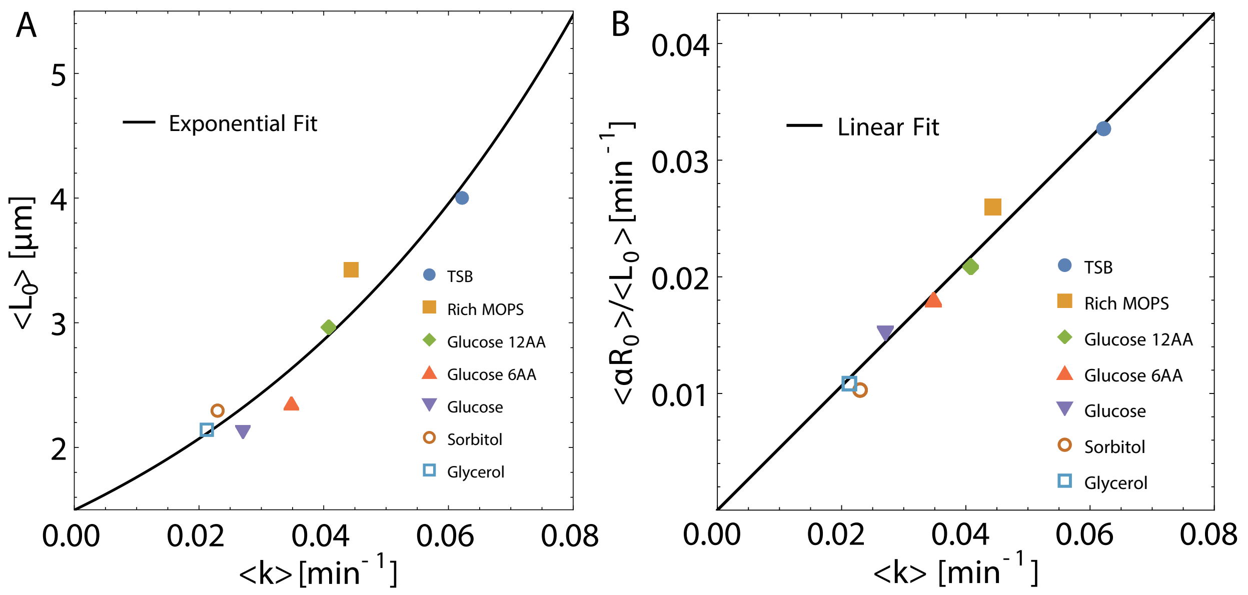


**Figure S 3. Cell length and ribosome abundance vs growth rate.** (A) Population-averaged initial cell length $\langle L_0 \rangle$ vs $\langle k \rangle$. The black curve shows an exponential fit to the data ($\langle L_0 \rangle = (1.50\ \mu m)\exp((16.19\ \text{min})\langle k \rangle)$). (B) Dependence of $\langle \alpha R_0 \rangle / \langle L_0 \rangle$ on elongation rate $\langle k \rangle$. The black line shows a linear fit to the data ($\langle \alpha R_0 \rangle / \langle L_0 \rangle = (0.56\ \mu m)\langle k \rangle$). Data for (A) and (B) taken from [4].

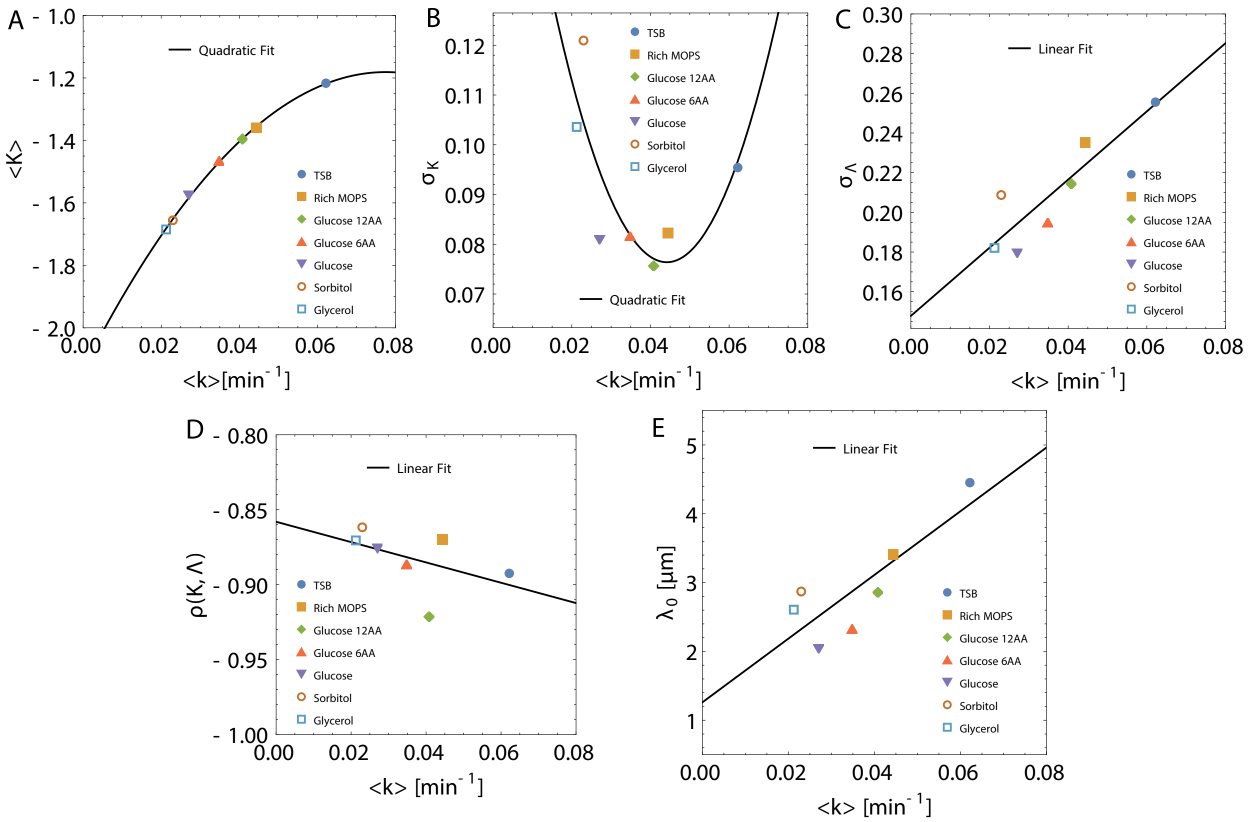


**Figure S 4. Growth-rate dependence of the parameters for the joint distribution $P(k,\lambda)$.** (A) $\langle K\rangle$ vs $\langle k\rangle$ from data with a quadratic fit ($(-158.15\,\mathrm{min}^{-2})\langle k\rangle^2 + (24.52\,\mathrm{min}^{-1})\langle k\rangle - 2.13$). (B) $\sigma_K$ vs $\langle k\rangle$ from data with a quadratic fit ($(62.42\,\mathrm{min}^{-2})\langle k\rangle^2 + (-5.51\,\mathrm{min}^{-1})\langle k\rangle + 0.20$). (C) $\sigma_\Lambda$ vs $\langle k\rangle$ from data with a linear fit ($(1.72\,\mathrm{min}^{-1})\langle k\rangle + 1.15$). (D) Correlation $\rho(K,\Lambda)$ vs $\langle k\rangle$ from data with a linear fit ($(-0.68\,\mathrm{min}^{-1})\langle k\rangle - 0.86$). (E) $\lambda_0$ vs $\langle k\rangle$ from data with a linear fit ($(46.3\,\mu m\,\mathrm{min}^{-1})\langle k\rangle + 1.26\,\mu m$). Data for (A-E) taken from [4].

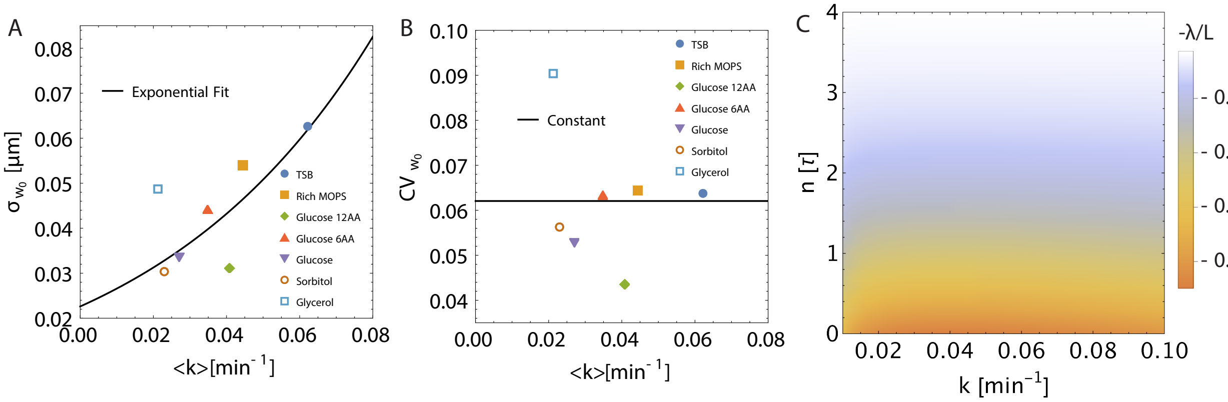


**Figure S 5. Properties of inter- and intra-gernerational width dynamics.** (A) $\sigma_{w_0}$ vs $\langle k \rangle$ from data with an exponential fit ($\sigma_{w_0} = 0.06\langle w_0 \rangle = (0.02\ \mu m)\exp((16.19\ \text{min})\langle k \rangle)$). (B) The coefficient of variation for $w_0$ is approximately constant based on prior fitting ($CV_{w_0} = \mu_{w_0}/\sigma_{w_0} = 0.06$). Data for (A) and (B) taken from [4]. (C) A colormap showing the approach of (13) to exponential growth ($-\lambda/L$) as a function of elongation rate $k$ and time $n$ in units of average interdivision time $\tau(k)$.

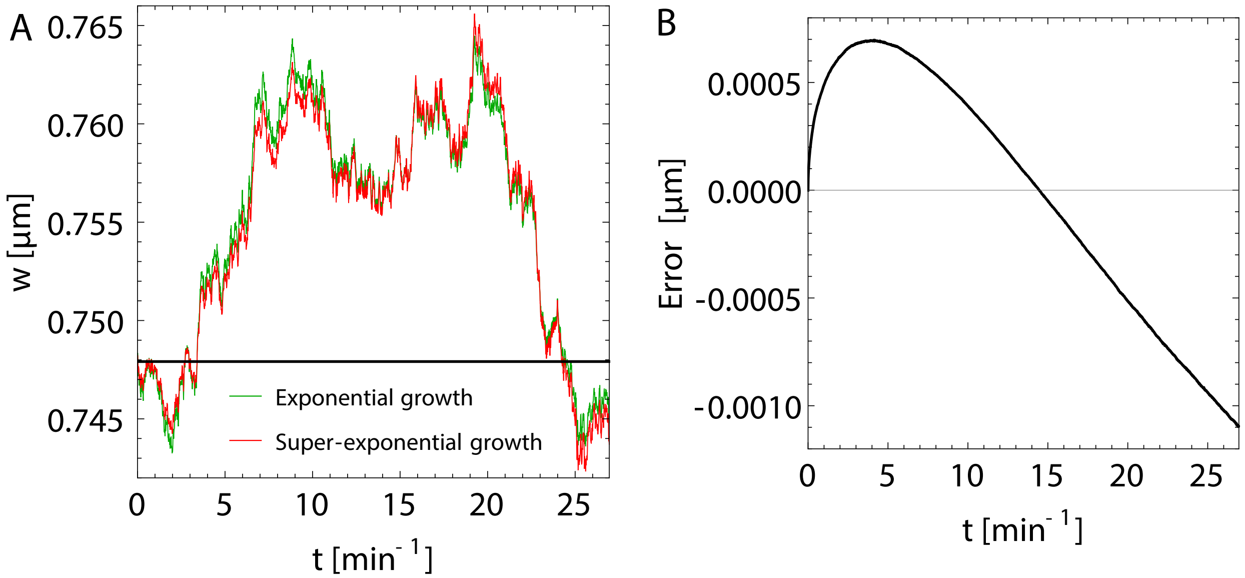


**Figure S 6. Super-exponential and exponential length growth produce similar width dynamics.** (A) Sample single cell width trajectories for exponential growth in green and super-exponential growth in red at $\langle k \rangle = 0.04\ \text{min}^{-1}$. Both trajectories start from the same initial condition and follow the same noise history with free parameters fit to experimental data as discussed in the main text. (B) Error between the exponential and super-exponential width SDE integrations as a function of time at $\langle k \rangle = 0.04\ \text{min}^{-1}$, averaged across 1000 cells. The qualitative shape and quantitative comparison to the absolute width values is seen across growth conditions.

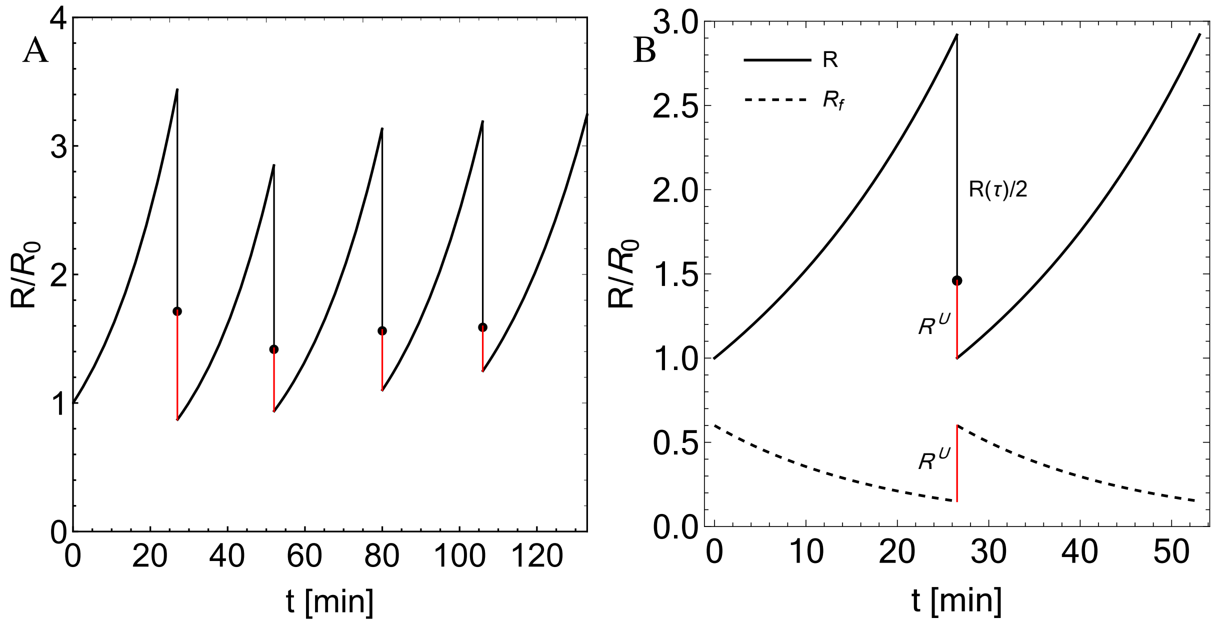


**Figure S** 7. **Inferred dynamics of ribosome abundance in *E. coli* in Glucose 12AA.** (A) Representative subsequent generations of ribosome dynamics inferred from experimental data of cell length. Black lines depict active ribosome abundance (normalized by the ribosome abundance at birth in the first generation), and red lines indicate the amount of ribosomes removed from the active pool following division ($R^{\mathrm{U}}$), calculated from the constants $k$ and $\alpha$ as described in the Appendix. (B) Ensemble-averaged data for ribosome abundance inferred from cell length data. Solid black depicts normalized active ribosome count while red highlights the average removed from the active pool following division ($R^{\mathrm{U}}$). Dashed lines show prediction of a potential model for ribosome removal from the active pool ($R_{\mathrm{U}}$) and allocated to a free ribosome pool ($R_{\mathrm{f}}$). See Appendix for model details. Data for (A) and (B) taken from [4].

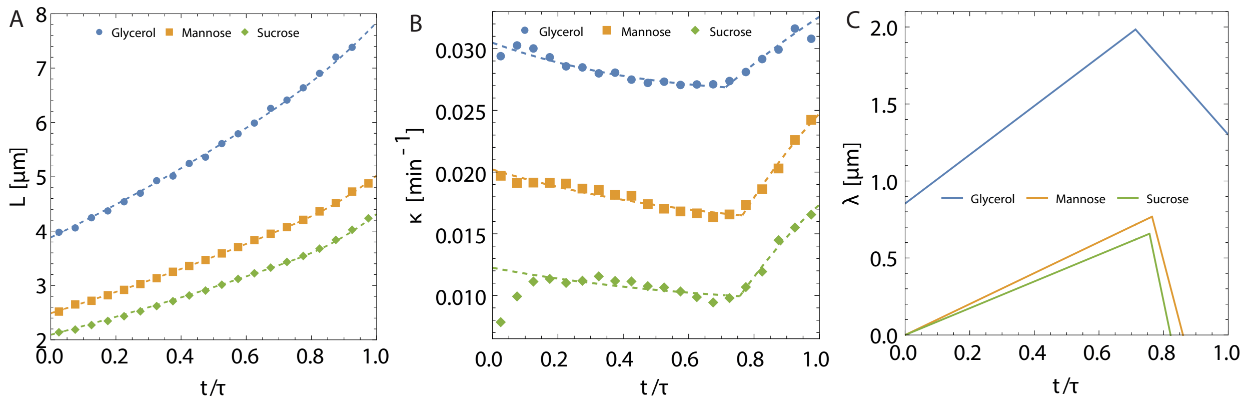


**Figure S** 8. **Modeling growth dynamics of *Bacillus subtilis*.** (A) Ensemble-averaged length of *B. subtilis* cells grown in three conditions at $37°C$ vs normalized time $t/\tau$, where $\tau$ is cell cycle duration. The average data is fit with the non-uniform growth model (2) with $\lambda$ piecewise linear (and continuous, see (C)) around a crossover timepoint. Error bars are negligible. (B) Ensemble-averaged instantaneous growth rate of *B. subtilis* cells grown in three conditions at $37°C$ vs normalized time $t/\tau$, where $\tau$ is cell cycle duration. Error bars show $\pm 1$ standard error in the mean. We show fits of (2) as found simultaneously with the length data in (A). (C) Resulting dynamics of $\lambda$ from the simultaneous fitting of model (2) to length and growth rate. Data for (A-C) taken from [39].
